# Targeted transcript quantification in single disseminated cancer cells after whole transcriptome amplification

**DOI:** 10.1101/616839

**Authors:** Franziska C. Durst, Ana Grujovic, Iris Ganser, Martin Hoffmann, Peter Ugocsai, Christoph A. Klein, Zbigniew T. Czyż

**Affiliations:** Experimental Medicine and Therapy Research, University of Regensburg, Regensburg, Germany; Fraunhofer-Institut für Toxikologie und Experimentelle Medizin, Project Group Personalized Tumor Therapy, Regensburg; Caritas-Krankenhaus St. Josef, Klinik für Frauenheilkunde, University of Regensburg, Germany

## Abstract

Gene expression analysis of rare or heterogeneous cell populations such as disseminated cancer cells (DCCs) requires a sensitive method allowing reliable analysis of single cells. Therefore, we developed and explored the feasibility of a quantitative PCR (qPCR) assay to analyze single-cell cDNA pre-amplified using a previously established whole transcriptome amplification (WTA) protocol. We carefully selected and optimized multiple steps of the protocol, e.g. re-amplification of WTA products, quantification of amplified cDNA yields and final qPCR quantification, to identify the most reliable and accurate workflow for quantitation of gene expression of the *ERBB2* gene in DCCs. We found that absolute quantification outperforms relative quantification. We then validated the performance of our method on single cells of established breast cancer cell lines displaying distinct levels of HER2 protein. The different protein levels were faithfully reflected by transcript expression across the tested cell lines thereby proving the accuracy of our approach. Finally, we applied our method on patient-derived breast cancer DCCs. Here, we were able to measure *ERBB2* expression levels in all HER2-positive DCCs. In addition, we could detect *ERBB2* transcript expression even in HER2-negative DCCs, suggesting post-transcriptional mechanisms of HER2 loss in anti-HER2-treated DCCs. In summary, we developed a reliable single-cell qPCR assay applicable to measure distinct levels of *ERBB2* in DCCs.

## Introduction

The analysis of systemically spread cancer via detection of disseminated cancer cells (DCCs) or circulating tumor cells (CTCs) in distant organs or blood, respectively, faces several technical challenges. First, the frequency of DCCs or CTCs is very low, e.g. ~two DCCs and ~one CTC can be found among 10^6^ nucleated cells in bone marrow and peripheral blood, respectively (1, 2), in breast cancer depending on the clinical stage. Second, micrometastatic cancer cells exhibit phenotypical and genetic heterogeneity affecting their malignant potential and susceptibility to therapy (3). Therefore, analysis of metastasis specifically at its early stages necessitates highly reliable methods enabling the analysis of single cells. Single-cell transcriptomes underlie dynamic changes that reflect functional and differentiation processes occurring in individual cells. Therefore, analysis of individual cell transcriptomes provides a first insight into cell functions relevant for disease progression or therapy resistance.

A single cell is believed to contain 1 pg of mRNA comprising transcripts expressed over several orders of magnitude, with the majority of genes being represented by less than 100 mRNA copies per cell (4). For the accurate assessment of heterogeneity among single cells, the applied workflows have to fulfill several specific requirements. First, a method dedicated for analysis of rare and unique cells should optimally provide sufficient amount of material to run all required downstream analyses. Second, the amplification of single-cell mRNA must be as accurate and comprehensive as possible to essentially preserve the qualitative and quantitative complexity of the sample. False-negative (technical drop-outs) and false-positive results should be reduced to a minimum. Third, an optimal workflow should be highly sensitive allowing detection of genes expressed at low levels. Various single-cell whole transcriptome amplification (WTA) methods have been developed (4, 5) permitting different types of downstream analyses utilizing qPCR (6–8), microarrays (9–11) or next generation sequencing (NGS) (12–15) as read-outs. Each of the available WTA technologies displays unique strengths and weaknesses reflected by differences in detection sensitivity (13, 14). Notably, available WTA strategies do not always provide an overview over the full-length transcripts of a single cell but instead are biased towards specific loci (7, 16) or the ends of the RNA molecules (17–19) limiting downstream analyses to either quantification of selected genes only (targeted analysis) or counting of mRNA molecules, thus limiting the ability to assess transcript diversity or mutational state analysis. The selection of an optimal technique for gene expression analysis in single cells depends on the specific research question, number of samples and genes to be analyzed, as well as the required sensitivity and costs. In the last years, powerful technological advances have been made in the area of single-cell RNA sequencing (scRNA-Seq), resulting in the development of technologies allowing the quantification of gene expression levels in single cells at much higher throughput (20). Progress in NGS and droplet microfluidics resulted in the development of methods enabling the analysis of hundreds to thousands of individual cells in a parallelized fashion (15) and detection of thousands of transcripts in a single experiment (21–23). However, high-throughput methodologies exhibit relative low mRNA capture rates (seizing only ~10-15% of transcripts expressed by each cell (18, 24)) that render them unsuitable for comprehensive analysis of rare cells. Moreover, NGS-based technologies require utilization of expensive hardware and generate high running costs. Importantly, scRNA-Seq workflows necessitate the use of advanced bioinformatic workflows and scripts to analyze the output data. Thus, users of such methods must have the knowledge and experience how to use (and in some cases establish) bioinformatic pipelines. In contrast, data analysis for qPCR such as relative and absolute quantification methods (25, 26), are well established, standardized and do not require a high level of expertise (27). Moreover, protocols for preparing qPCR samples are simpler and result in higher sensitivity and reproducibility as compared to NGS-based approaches (7, 27). Importantly, single-cell qPCR workflows exhibit high levels of reliability and wide and dynamic detection ranges, making them exceptionally well-suited for targeted gene expression analyses in singles cells, where sensitivity is essential and the amount of target genes is low (7, 27). Therefore, the present study aimed to develop a single-cell qPCR assay to quantify gene expression changes in single cells, specifically in patient-derived DCCs. We established a workflow comprised of single-cell WTA, re-amplification of single-cell cDNA, post-WTA normalization of cDNA quantities and qPCR-based data analysis. The new assay provides means for measuring expression levels of individual pre-selected genes in WTA products generated from single cells in an accurate and reliable fashion.

## Materials and methods

### Cell lines

BT-474 (ACC 64) and MCF-7 (ACC 115) breast cancer cell lines were obtained from German Collection of Microorganisms and Cell Cultures (DSMZ). MCF-10A (CRL-10317), a non-tumorigenic mammary epithelial cell line was obtained from American Type Culture Collection (ATCC). ZR-75-1 (CRL1500, ATCC) and MDA-MB-453 (ACC 65, DSMZ) cells were purchased from DSMZ. The identity of all cell lines was confirmed by DNA finger printing analysis utilizing the GenePrint® 10 System (Promega). BT-474 and MDA-MB-453 cells were cultivated in DMEM medium (Pan-Biotech) supplemented with 10% FCS (Sigma-Aldrich), 2 mM L-Glutamine (Pan-Biotech) and 1% Penicillin/Streptomycin (Pan-Biotech). ZR-75-1 and MCF-7 cells were propagated in RPMI 1640 medium (Pan-Biotech) supplemented with 10% FCS, 2 mM L-Glutamine and 1% Penicillin/Streptomycin. In addition to other components, medium for MCF-7 cells contained 1 mM Sodium-Pyruvate (Sigma-Aldrich). MCF-10A cells were cultured in DMEM/F12 medium (Pan-Biotech) supplemented with 5% Horse serum (Sigma-Aldrich), 1% Penicillin/Streptomycin, 20 ng/ml EGF (Sigma-Aldrich), 0.5 μg/ml Hydrocortisone (Sigma-Aldrich), 10 μg/ml Insulin (Sigma-Aldrich) and 0.1 μg/ml Cholera toxin (Sigma-Aldrich).

### Flow cytometry

Cell line cells were harvested using 0.05% Trypsin/EDTA (Pan-Biotech) for 3 min and stained using mouse anti-human HER2 antibody conjugated to FITC (20 μg/ml, clone 24D2, BioLegend). Cells were incubated with anti-HER2 antibody for 20 min at 4°C in the dark and washed once with PBS supplemented with 1% FCS prior to the FACS analysis. A corresponding isotype control (Mouse IgG1, κ-FITC labeled, eBioscience) was used in every staining experiment to determine the background level of fluorescence and set the threshold for specific staining signals. Cells were analyzed on a BD FACS Canto II instrument (BD Bioscience) equipped with FACS DIVA v7.0 software (BD Bioscience). Sorting of single cells was performed with a FACSAria cell sorter (BD Bioscience). FlowJo v10 (Treestar) software was used for analysis of the obtained data.

### Patient sample

Pleural effusion from a patient with HER2-positive metastatic breast cancer was obtained from the Caritas Hospital St. Josef in Regensburg. The ethics committees of the University of Regensburg approved the sampling and genetic analysis of the isolated cells (ethics vote number 17-672-101). The donor provided written informed consent for processing of the clinical material. The patient received her regular dose of trastuzumab / pertuzumab early in the morning and the pleural effusion was sampled in the late afternoon. The sample was centrifuged for 10 min at 300×*g* and cells were washed once using Hank’s solution (Biochrom). 2×10^6^ cells were double-stained using anti-EpCAM (1:50, clone HEA-125, Miltenyi) and anti-HER2 antibodies directly conjugated to PE and FITC, respectively. After 15 min incubation at 4°C on a roller mixer in the dark, cells were washed once with 1× PBS. EpCAM staining was used to detect DCCs. Stained sample was screened on an Olympus IX-81 inverted fluorescent microscope (Olympus) for the presence of marker positive cells. Single cells were manually isolated with a micromanipulator (Eppendorf, PatchMan NP2) as previously described (9). Isolated cells were picked in 1 μl of 1× PBS and transferred into 4.4 μl of lysing buffer containing 4 μl mTRAP™ Lysis Buffer (Active Motif) and 0.4 μl (10 ng) tRNA from E. coli MRE 600 (Roche). Picked cells suspended in the lysis were stored at −80°C until further processing, i.e. WTA.

### Whole Transcriptome Amplification (WTA) of single cells and quality control

Single cells were subjected to WTA using a previously described protocol (9). Primer sequences for WTA and re-amplification are provided in S2 Table. Quality of isolated primary cDNA amplification products and re-amplified WTA products was examined utilizing endpoint PCR controlling for expression of three housekeeping genes: *EEF1A1, ACTB* and *GAPDH*. Each PCR reaction was composed of the following ingredients: 1 μl 10× FastStart PCR Buffer comprising 20 mM MgCl2 (Roche Diagnostics), 0.2 μl dNTPs (10 mM each), 1 μl of Primer-Mix (8 μM of each forward and reverse primer for each tested gene – *EEF1A1*, *ACTB* and *GAPDH*; S1 Table), 0.2 μl BSA (20 mg/ml, Roche Diagnostics), 6.5 μl HPLC Gradient Grade H_2_O, 0.1 μl FastStart Taq Polymerase (5 U/μl, Roche Diagnostics) and 1 μl of fivefold diluted primary WTA product. PCR was performed on MJ Research Peltier Thermal Cycler Tetrad (Bio-Rad) using the following program: initial denaturation at 95°C for 4 min was followed by 32 cycles of 30 s at 95°C, 30 s at 58°C, 90 s at 72°C and a final elongation step of 7 min at 72°C. Positive and negative controls were included in every PCR run.

### Gene expression analysis using endpoint PCR

Endpoint PCR was conducted to verify the presence of selected genes of interest (S1 Table) in WTA products. Primary or re-amplified WTA product was diluted five times with water and used as template in each test PCR (1 μl per sample). Templates were mixed with 1 μl of 10× PCR buffer containing PCR Grade dNTP Mix (10 mM each), 0.5 μl of forward and reverse primer (8 μM each; S1 Table), 0.25 μl BSA (10 mg/ml, Roche Diagnostics), 0.1 μl PANScript DNA Polymerase (5 U/μl, Pan Biotech) and HPLC Grade H_2_O to a final volume of 10 μl. PCR was run using the following program: initial denaturation of 4 min at 95°C was followed by 42 cycles of 30 s at 95°C, 30 s at 58°C, 90 s at 72°C and a final elongation step for 7 min at 72°C. Positive and negative control was included in every PCR run.

### Re-amplification of primary WTA single-cell products

Re-amplification of primary WTA products was performed in a reaction volume of 50 μl comprising 5 μl Expand Long Template Buffer 1 (Expand Long Template PCR System, Roche Diagnostics), 6 μl of CP2-15C or CP2-9C primer (2.88 μM; S2 Table), 1.75 μl dNTPs (10 mM each), 7.5 μl 20% Formamide (Sigma Aldrich), 1.5 μl DNA Pol Mix (5 U/μl, Expand Long Template PCR System, Roche Diagnostics), 27.25 μl PCR HPLC Gradient Grade H_2_O and 1 μl template (primary WTA product). Re-amplification was run on PTC DNA Engine 2 Tetrad Thermocycler using the following program: 1 min at 95°C, 5 cycles comprising 15 s at 94°C, 1 min at 60°C and 3 min 30 s at 65°C, 3 cycles of 15 s at 94°C, 1 min at 60°C and 3 min 30 s at 65°C (elongation step was extended by 10 s per cycle) and a final elongation step of 7 min at 65°C. Negative control was included in every run. Quality of re-amplified product was examined using the same endpoint PCR assay as the one used for primary WTA products (see above).

### Quantitative Real-Time PCR and statistical methods

Quantitative PCR (qPCR) was performed for selected genes of interest (*ERBB2* and reference genes) using a LightCycler 480 instrument (Roche). Each qPCR comprised 5 μl of the template cDNA, 10 μl iQTM SYBR® Green Supermix (Bio-Rad), 1 μl of each forward and reverse primer (8 μM each, S1 Table) and 3 μl PCR HPLC Gradient Grade H_2_O. Quantitative PCR was run using the following program: 1 cycle for 5 min at 95°C (temp. ramp of 4.4°C/s), 38 cycles for 20 s at 95°C (ramp 4.4°C/s), 15 s at 58°C (ramp 2.2°C/s) and 15 s at 72°C (ramp 4.4°C/s; fluorescence signal was measured at the end of each elongation step). Subsequently, melting curves were generated using the following procedure: 1 cycle for 5 s at 95°C (ramp 4.4°C/s), 1 cycle for 1 min at 50°C (ramp 2.2°C/s), 1 cycle of DNA melting, wherein temperature was continuously increased to 95°C with a ramp of 0.11°C/s with continuous fluorescence measurement 5 times/s) followed by a final cooling to 40°C for 30 s (2.2°C/s). Melting plots were examined to validate the specificity of PCR amplification. Samples with appearance of melting curves different than expected were excluded from downstream analyses. Crossing point (Cp) values were determined with the LightCycler 480 Software using the second derivative maximum method applying the high sensitivity algorithm. All single-cell WTA products were analyzed in technical triplicates. Cp-values were averaged across the technical replicates before further data processing. Samples with average Cp-values >33 were considered as negative. No template control was included in any qPCR run.

### Relative qPCR quantification analysis

Template cDNA (i.e. WTA or re-amplified WTA) was diluted 50 times and 5 μl of the diluted cDNA was used per qPCR reaction. Primer efficiencies were calculated based on the analysis of standard curves generated using serially diluted WTA samples. For each measured sample, the mean Cp-value of the gene of interest was normalized to the expression of a single reference gene by calculating ΔΔCp values according to the formula as described before (25). Visualization of relative qPCR quantification data and downstream analyses were conducted utilizing log2-transformed ratios resulting in negative ΔΔCp values.

### Absolute qPCR quantification analysis

Absolute qPCR quantification analysis of *ERBB2* expression levels necessitates the normalization of the template input. Quantification of cDNA yields in the individual samples was spectrophotometrically conducted using either the NanoDrop 2000 instrument or the Qubit dsDNA BR kit and the Qubit fluorometer. To allow an accurate measurement of cDNA, yields of WTA products were either purified or subjected to double-stranded DNA reconstitution (see below). DNA input for each qPCR was normalized to 5 ng and run as described above. The obtained Cp-values generated by the LightCycler 480 Software (Roche) were converted into log10 copy numbers. For this, a standard curve measurement was conducted with serially diluted cDNA standards comprising amplicons of the *ERBB2* transcript. Concentrations of the standards ranged from 1.00E-08 to 1.00E-04 ng/μl covering copy numbers from 3.00E02 to 3.00E06 molecules/μl.

### Purification of WTA products

Purification of WTA samples was conducted to remove buffer and remaining WTA reagents (dNTPs, proteins and primers) that may negatively influence downstream processes (i.e. cDNA yield quantification and qPCR). 10-15 μl of primary or re-amplified WTA product was purified using QIAquick PCR Purification Kit (QIAgen) according to the manufacturer’s instruction with several changes: (i) No pH-indicator was added to the PB buffer. (ii) Purified cDNA was eluted from the purification column using HPLC Gradient Grade H_2_O instead of EB buffer provided by the manufacturer of the kit. (iii) PCR HPLC Gradient Grade H_2_O (typically 15-20 μl) used for elusion was pipetted on the silica membrane of the column, followed by 5 min incubation at room temperature prior to the final centrifugation (elution) step. To allow a more optimal distribution of elution liquid in the silica membrane on the purification column, spin assembly was centrifuged at 500 rpm for 30 s followed by the final centrifugation at 13,000 rpm for 60 s. Concentration of each purified sample was measured using NanoDrop 2000c (Thermo Fisher Scientific) utilizing 1 μl of purified cDNA.

### Double stranded DNA reconstitution of WTA products

Double strand cDNA (dscDNA) reconstitution was conducted to convert single stranded cDNA amplicons present in the WTA products into double stranded molecules allowing fluorometric quantification of cDNA yields without the need for prior purification of the WTA samples. dscDNA-synthesis was performed with 10 μl of the template DNA (i.e. primary or re-amplified WTA products) added to 1 μl Expand Long Template Buffer 2 (Expand Long Template PCR System, Roche Diagnostics), 1 μl dNTPs (10 mM each), 1 μl CP2-15C (2 μM; S2 Table), 0.5 μl DNA Pol Mix (Roche Diagnostics) and 6.5 μl HPLC Gradient Grade H_2_O. Reactions were run using a PTC DNA Engine 2 Tetrad Thermocycler for 2 h at 68°C. Concentration of resulting dscDNA was measured using Qubit BR Kit and Qubit 2.0 Fluorometer (Thermo Fisher Scientific) according to the manufacturer’s instruction using the Broad Range Assay. 2 μl of dscDNA were used for each measurement.

### Bioanalyzer assay

To analyze the fragment size distribution of WTA samples, Agilent High Sensitivity DNA Kit (Agilent Technologies) was used according to the manufacturer’s instructions. WTA samples underwent dscDNA reconstitution to determine the concentration of dscDNA. Samples were further diluted to 1 ng/μl and 1 μl was used for analysis in the Agilent 2100 Bioanalyzer.

### Statistical analysis

Statistical analysis was conducted using the GraphPad Prism 6.01 software package (GraphPad Software Inc). −ΔΔCp values and log10 copy number values were tested for Gaussian distribution using the D’Agostino and Pearson omnibus normality tests. Pearson’s (parametric) or Spearman’s (non-parametric) correlation tests were used to compare sample sets. The unpaired t-test (two-tailed, CI=95%) with Welch’s correction was used to determine the statistical significance of differences between gene expression levels measured in BT-474 and MCF-7 cells. One-way ANOVA test with Tukey’s multiple comparisons test was used to assess the statistical significance between gene expression in more than two tested cell populations. Values of p<0.05 (marked with a * in the figures) were considered statistically significant, whereas p<0.01 (**), p<0.001 (***) and p<0.0001 (****) were recognized as highly significant.

## Results

### Relative quantification of gene expression in single cells

Relative transcript quantification by qPCR, for which the expression level of a gene of interest is calculated relative to the expression of one or multiple reference gene(s), so called housekeeping genes, is widely used to measure mRNA levels in multiple sample types (28). However, to assure maximal accuracy of measurements one has to select reference genes that (i) are stably and consistently expressed at detectable levels across the studied sample collective and (ii) remain unaltered in the experimental set-up used. To identify the most suitable reference genes, we assembled a shortlist of putative housekeeping genes using three different approaches. First, we selected eight genes (*POLR2A*, *G6PD*, *HPRT1, ABCF1, GPS1, VPS72, CANX* and *TBP*) involved in basic metabolic processes as we hypothesized that these need to be expressed in every cell. Second, we re-evaluated a series of single-cell microarray-based gene expression profiling experiments to find most consistently and stably expressed genes. For this, we used samples stemming from various experimental conditions including *ex vivo* isolated cell types that had been amplified by our previously reported WTA method prior to microarray hybridization (9, 29–31). Utilizing this approach, we identified 1,544 candidate genes. From this group we selected seven candidate genes (*WNT10A*, *IRX3*, *MYOM1*, *NUDT13*, *ASCL2*, *EHMT2* and *DUSP15*) exhibiting strong and constant expression for further testing. Lastly, we picked additional five candidates (*EMC7*, *RAB7A*, *REEP5*, *VCP* and *PSMB4*) from a previously reported RNA-Seq-derived list of highly uniformly expressed housekeeping genes (32). We designed and tested one primer pair for every candidate gene (S1 Table). Following initial tests, eight genes (*ABCF1*, *GPS1*, *VPS72*, *WNT10A*, *IRX3*, *MYOM1*, *NUDT13*, *ASCL2*) were excluded due to unspecific amplification (multiple amplification products and/or other than expected RFLP patterns in endpoint PCR; data not shown). The remaining candidate genes (*POLR2A*, *G6PD*, *HPRT1, CANX*, *TBP*, *EHMT2*, *DUSP15*, *EMC7*, *RAB7A*, *REEP5*, *VCP* and *PSMB4*) were then examined for stability and uniformity of expression in a cohort of WTA products obtained from cell pools and single cells isolated from various tissues: (i) fourteen cell pools (consisting of approximately 40 cells each) from patients with acute lymphocytic leukemia, (ii) sixteen single EpCAM+ DCCs from patients with prostate cancer and (iii) fifteen cell pools (consisting of a few hundred cells) from primary cultures of patients with melanoma, bronchial or prostate carcinoma (five cell pools each). First screenings were conducted using endpoint PCRs. Surprisingly, none of the twelve genes under investigation was consistently expressed in all of the tested samples. The transcript detection rate varied considerably (11-93%) between different candidate genes (Fig 1A, S3 Table). To exclude low-abundantly expressed genes, we set a detection rate cut-off to 60% (i.e. the tested gene must be expressed in more than 60% of samples) resulting in omission of three rarely expressed genes (*CANX*, *DUSP15* and *TBP*) from further analyses (Fig 1A). Next, we re-analyzed the same sample set by qPCR to assess the level and uniformity of the remaining nine reference genes. Melt curve analyses revealed unspecific amplification (diverging appearance of melt curves across samples) of the *PSBM4* gene resulting in its exclusion from further analyses. *RAB7A*, *EMC7* and *REEP5* genes showed the most consistent expression across all tested samples (Fig 1B). To assess the uniformity of expression, we calculated log2-transformed ratios (−ΔΔCp values) (see Materials and methods) for each putative reference gene against all remaining ones. We hypothesized that stably expressed genes exhibit low variances of −ΔΔCp. Among the genes tested, *POLR2A* showed the most stable expression (Fig 1C, S4 Table). Notably, for all tested genes except *RAB7A* the variance of measured −ΔΔCp was higher in single-cell WTA products as compared to those generated from cell pools (Fig 1C), indicating that the expression of individual genes is noisier at the single-cell level as compared to bulk measurements. Finally, taking into account gene expression consistency and uniformity, five genes (*RAB7A*, *EMC7*, *REEP5,*, *POLR2A* and *HPRT1*) were chosen as potential reference genes for further testing.

Next, we examined whether normalization to each of the selected reference genes enables robust quantitative measurement of mRNA levels of a selected target gene in single-cell WTA products. To this end, we investigated expression levels of the *ERBB2* gene in two breast cancer cell lines - BT-474 and MCF-7 - known to differ in expression levels of HER2, the protein product of the *ERBB2* gene (33, 34). FACS analysis revealed a 62-fold higher HER2 expression in BT-474 as compared to MCF-7 cells (Median FITC [BT-474] = 24,659; Median FITC [MCF-7] = 399; Fig 1D). From either cell line twenty-two single cells were picked by micromanipulation and subsequently subjected to WTA. All cells were tested for the presence of transcripts of three housekeeping genes (*ACTB*, *GAPDH* and *EEF1A1*; WTA quality control assay – see Material and methods), the target gene (*ERBB2*) and the five reference genes (*RAB7A*, *EMC7*, *REEP5, POLR2A* and *HPRT1*) using endpoint PCRs. Drop-out rates (i.e. failure to detect the expression of a tested transcript) for each PCR assay are documented in Table 1 and S5 Table. Of note, drop-outs were detected at a considerable frequency for transcripts of the tested reference genes, which hindered measurement of gene expression in single cells when using relative quantification. One MCF-7 (6.7 %) and six BT-474 single cells (30 %) had to be excluded from qPCR analysis due to non-detectable expression of reference transcripts in the endpoint PCR analysis (Table 1). Still, we tested the remaining fourteen single cells from each BT-474 and MCF-7 cell line by qPCR. Subsequent statistical analysis of single-cell qPCR data showed a significant difference in *ERBB2* transcript level between BT-474 (HER2^hi^) and MCF-7 (HER2^lo^) calculated for all target-reference gene pairs (Fig 1E). As a consequence of the pre-selection by endpoint PCR, no further drop-outs were observed for qPCR. In summary, we identified five reference genes suitable for relative quantification of mRNA expression levels in single-cell WTA products.

**Table 1.**
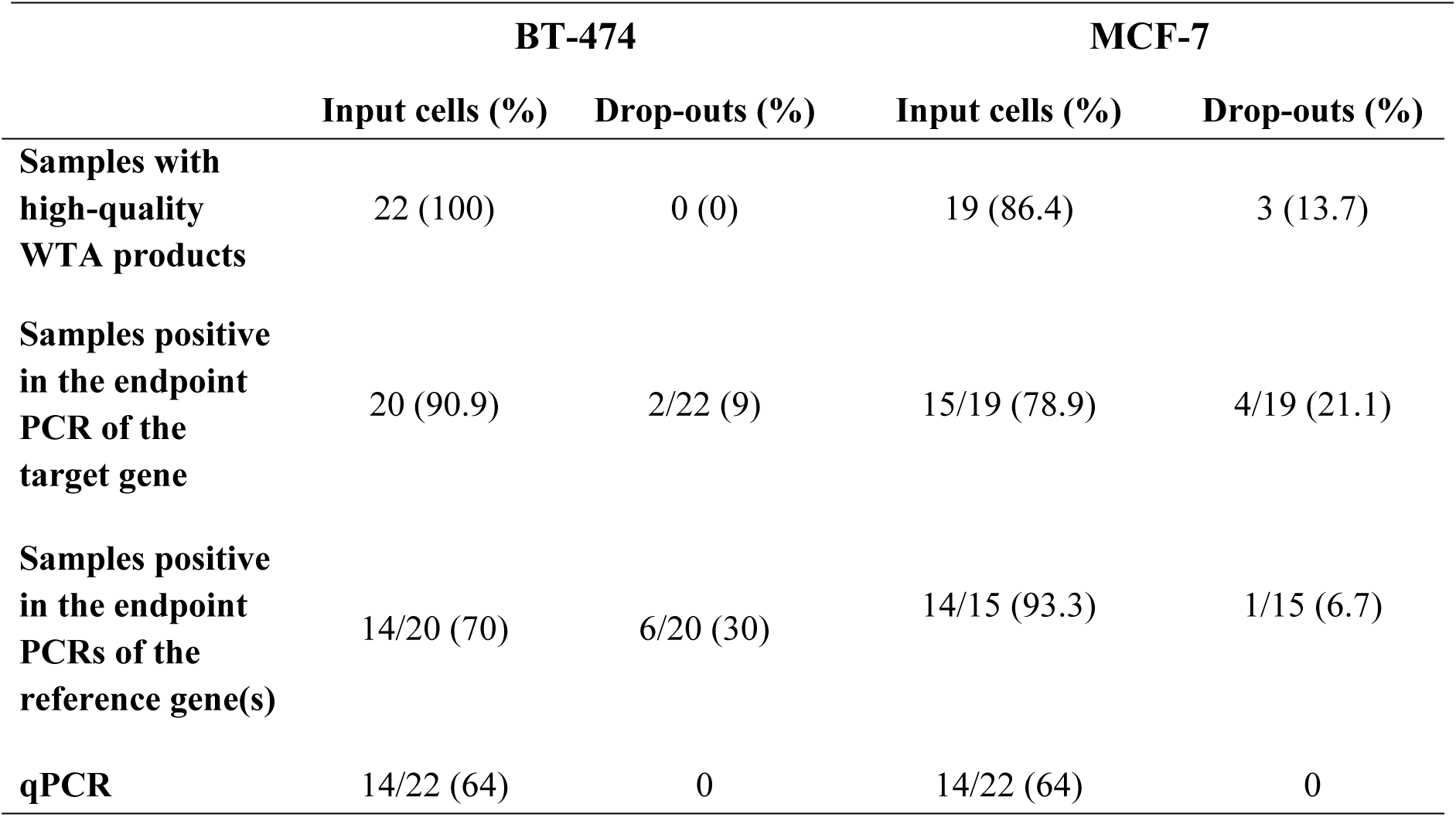
Stepwise drop-out rates during sample selection for qPCR.

**Fig 1.**
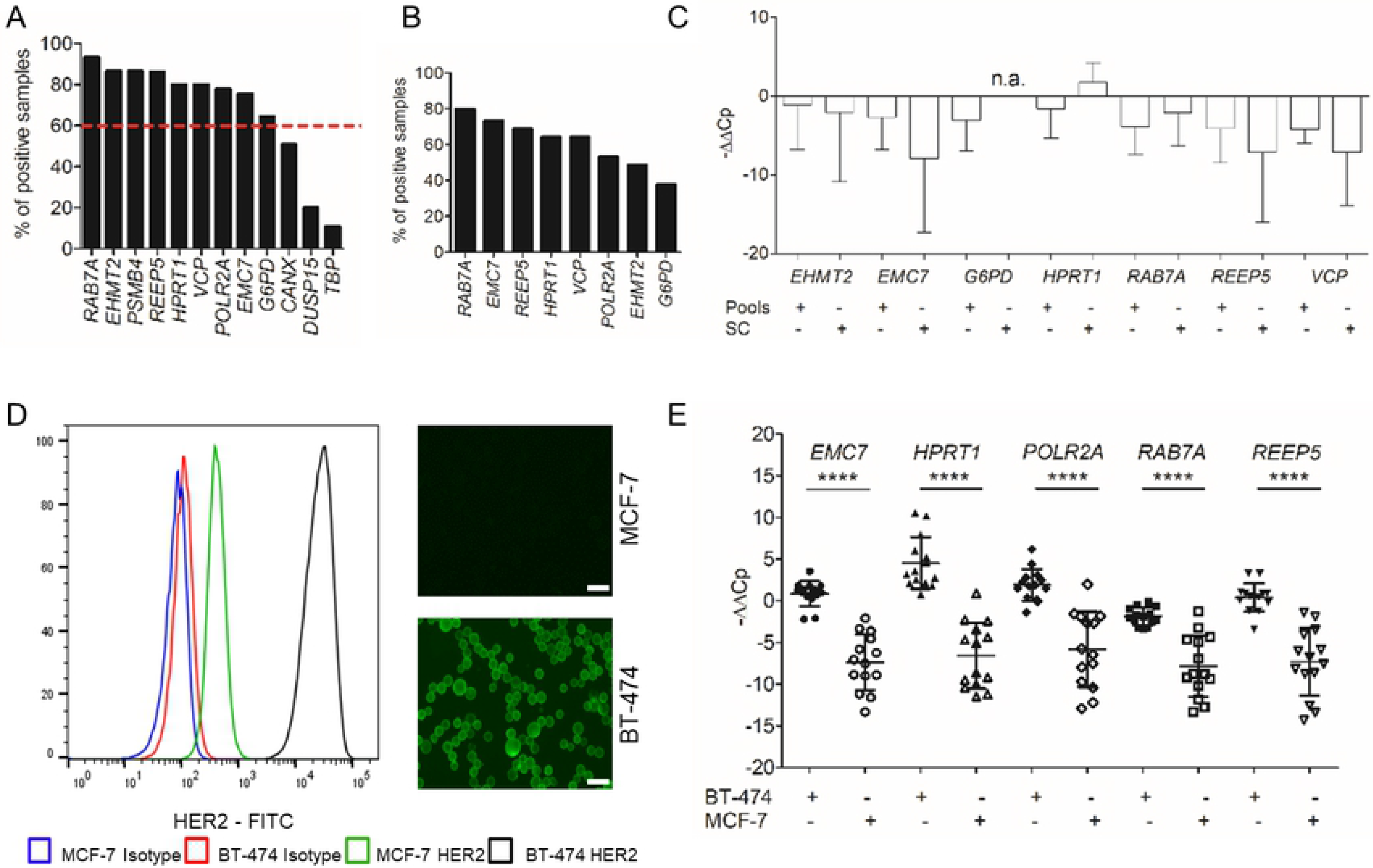
Relative qPCR quantification of single cell transcripts. (A) Stable and uniform gene expression (depicted as percentage of samples tested positive for selected transcripts) of twelve genes examined by endpoint PCRs of WTA products obtained from three sample sets including cell pools and single cells. The red dashed line indicates the threshold for further testing (60%). (B) Uniformity of gene expression (%) of the remaining eight candidate reference genes as measured by qPCR. (C) Stability of gene expression depicted as −ΔΔCp (log2-transformed ratios) calculated for *POLR2A* vs the remaining candidate genes. Expression of *G6PD* could not be measured in single cells (n.a. = not assessed). SC = single-cell WTA products, Pools = WTA products generated from a pool of cells. (D) HER2 expression in BT-474 and MCF-7 breast cancer cells analyzed by FACS (left panel) and microscopically (right panels). HER2 expression was 62-fold higher in BT-474 as compared to MCF-7 cells. Median FITC (BT-474) = 24,659; Median FITC (MCF-7) = 399. Microscope settings corrections: Brightness: +20%, contrast: −40%; Scale bar 20 μm. (E) Relative quantification of *ERBB2* expression at the single-cell level in BT-474 and MCF-7 cells calculated separately using single reference genes as indicated. −ΔΔCp were calculated and plotted for every single-cell qPCR measurement. Mean ± SD of −ΔΔCp values; Unpaired t-test with Welch’s correction; *** p<0.001, **** p<0.0001.

### Absolute quantification of gene expression in single cells

In contrast to relative qPCR quantification, absolute quantification does not require measurements of reference genes in tested samples but careful normalization of cDNA inputs. For this, concentration of template cDNA needs to be determined and equalized across all samples. However, since single-cell WTA products are mixtures of single stranded and double stranded cDNA amplicons (sscDNA and dscDNA, respectively) as well as primers, dNTPs and other reagents used in the WTA procedure, direct measurement of cDNA yields utilizing spectrophotometric or fluorometric methods is prone to error. We therefore compared two approaches to assess DNA concentrations in single-cell WTA products. In the first approach, samples were purified using the QIAquick PCR Purification Kit (to remove residual buffer, dNTPs and primers) prior to spectrophotometric measurement. In the second procedure, samples were subjected to dscDNA reconstitution of the WTA products enabling subsequent fluorometric measurement using dsDNA-binding dyes. Both protocols were applied to the WTA products of single BT-474 and MCF-7 cells mentioned above. Subsequently, 5 ng of the obtained cDNA were used in every qPCR to measure expression levels of the *ERBB2* gene. In line with results obtained by relative quantification, absolute quantification allowed us to detect distinct expression levels of the *ERBB2* gene in HER2^hi^ and HER2^lo^ cells using both cDNA processing strategies (Fig 2A and 2B). Of note, only one BT-474 single cell that underwent dscDNA reconstitution had to be excluded from qPCR analysis due to discrepant amplification plots (S5 Table). Accordingly, we can report a drop-out rate of only 2% (1/44) for single-cell samples subjected to absolute quantification (i.e. relative to both MCF-7 and BT-474 cells after WTA purification or dscDNA reconstitution). As an advantage of absolute quantification over relative quantification, calculated differences in *ERBB2* copy numbers more faithfully reflected the 62-fold difference in HER2 protein expression level between HER2^hi^ and HER2^lo^ cells (50-fold or 51-fold more *ERBB2* copy numbers in BT-474 cells as compared to MCF-7 cells using dscDNA or purified samples, respectively) (Fig 1D, Fig 2A and 2B). To better define the impact of sample pre-treatment (i.e. dscDNA synthesis or WTA purification) on the accuracy of qPCR measurements, we generated more data points not restricted to *ERBB2* alone by measuring the expression levels of all five previously selected reference genes (*RAB7A*, *EMC7*, *REEP5, POLR2A* and *HPRT1*) (S5 Table). Careful analysis of the qPCR data generated for *RAB7A* gene revealed that two samples (one generated from MCF-7 and one from BT-474) subjected to dscDNA exhibited discrepant amplification curves as compared to other samples included in the collective and were therefore excluded from further analyses (S5 Table). Direct comparison of calculated −ΔΔCp values obtained for all possible gene pairs showed a high level of correlation between results generated using unprocessed primary WTA products and results obtained after dscDNA reconstitution or WTA purification (Fig 2C and 2D). Here, −ΔΔCp values obtained from purified samples correlated better with the corresponding values generated from untreated specimens as compared to samples subjected to dscDNA reconstitution (Fig 2C and 2D). Based on this correlation analysis and the observed high drop-outs of samples subjected to dscDNA reconstitution, we concluded, that WTA purification is more suited as a cDNA pre-treatment method than dscDNA reconstitution. Summarizing, we established a workflow consisting of i) single-cell WTA of cells isolated by e.g. micromanipulation including an optional WTA re-amplification step (see below), ii) a WTA quality control assay (endpoint PCR targeting the three housekeeping genes *ACTB*, *GAPDH* and *EEF1A1*), iii) an optional endpoint PCR for the selected gene of interest (acting as screening to select samples suitable for downstream analyses), iv) purification of amplified cDNA, v) spectroscopic cDNA quantification, vi) normalization of cDNA input and vii) qPCR measurement with absolute quantification used for data analysis (Fig 2E). This workflow was utilized in the subsequent experiments.

**Fig 2.**
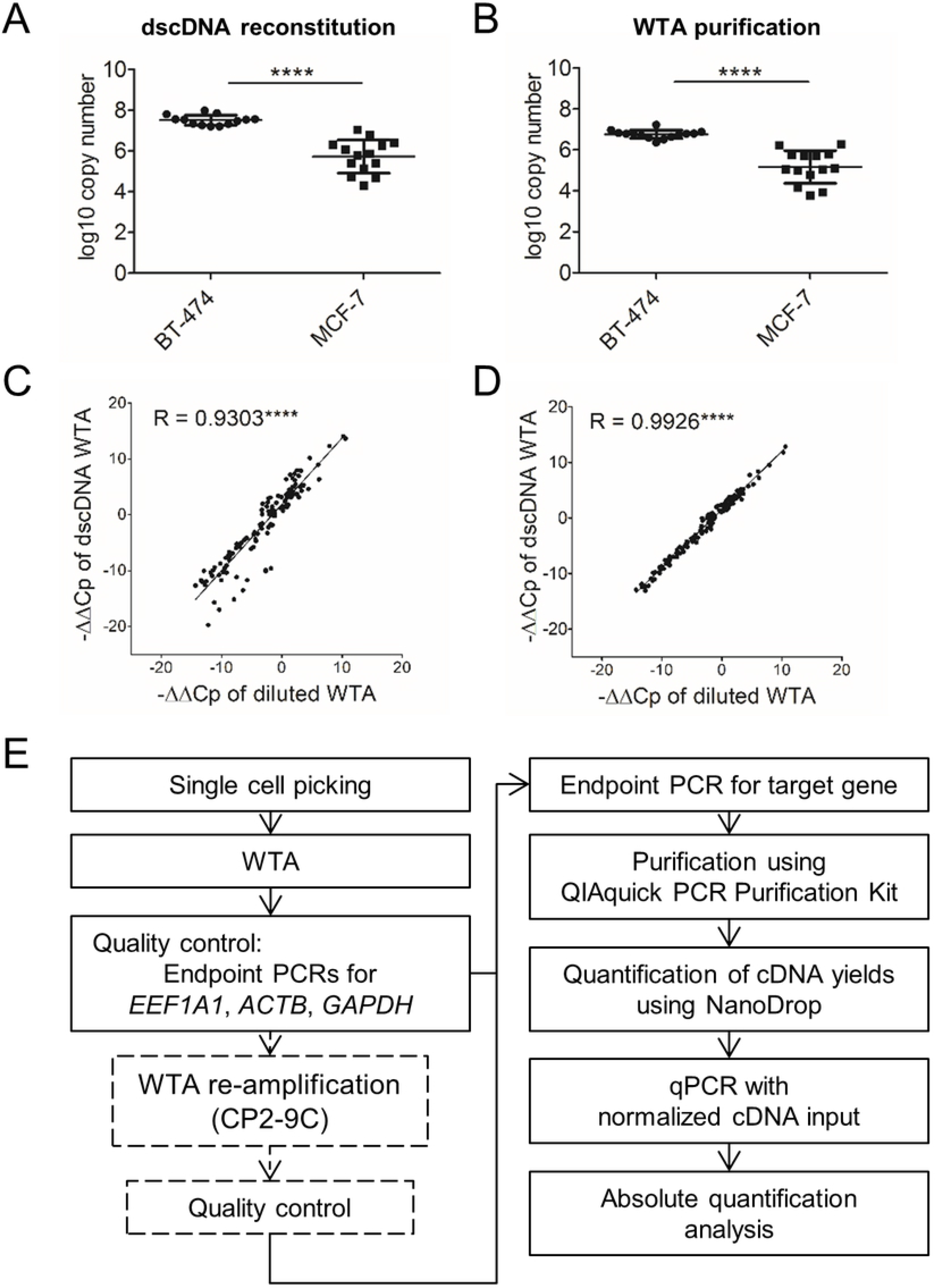
Novel qPCR-based workflow for highly accurate gene expression analysis in single cells. (A, B) Quantification of the *ERBB2* gene expression levels in MCF-7 and BT-474 single cells using absolute qPCR quantification. WTA products underwent either double-stranded cDNA (dscDNA) reconstitution (A) or purification using QIAquick PCR Purification Kit (B) prior to normalization of cDNA input and qPCR. Cp values were converted to log10 copy numbers using an external standard curve. Mean ± SD; Unpaired t-test with Welch’s correction; **** p<0.0001. (C, D) Correlation of *ERBB2* expression levels obtained using relative quantification strategy in diluted WTA samples as compared to WTA products subjected to dscDNA reconstitution (C) or purification (D). Each point represents a −ΔΔCp value calculated for one *ERBB2*-reference gene pair (−ΔΔCp for all target/reference gene combinations were plotted). Pearson’s correlation coefficient R. **** p<0.0001. (E) Workflow of the established qPCR assay for profiling gene expression levels in single cells using the absolute quantification strategy.

### Sensitivity of the new single-cell qPCR workflow

Dependency of relative quantification on a simultaneous expression of multiple genes (i.e. target and reference gene(s)) hinders quantification if expression of one of the tested transcripts cannot be detected. Missing expression values were prominent in single-cell samples before precluding gene expression analysis of these samples using when using the relative quantification method (Table 1, S5 Table). In contrast, absolute quantification provided the means for a more robust and reliable gene expression profiling in single-cell WTA products. Therefore, we decided to study the accuracy of the absolute quantification method in greater detail. We attempted to measure finer differences in gene expression levels than those quantified when analyzing BT-474 and MCF-7 cells (62-fold difference). Accordingly, we chose three well-established breast cancer cell lines (MCF-10A, ZR-75-1 and MDA-MB-453) exhibiting similar, partially overlapping, levels of HER2 protein expression as determined by flow cytometry (Fig 3A and 3B) (34). FACS analysis of ZR-75-1 showed a 3.5-fold higher expression level of HER2 protein compared to MCF-10A, and a 2.3-fold lower amount of HER2 protein compared to MDA-MB-453 cells (Fig 3B). We utilized FACS to isolate cell populations exhibiting distinct levels of HER2 (similar within a given cell population but distinct from other groups) from each cell line (Median FITC [MCF-10A] = 436; Median FITC [ZR-75-1] = 1,529; Median FITC [MDA-MB-453] = 3,575; Fig 3B). Subsequently, ten single cells from each sorted subpopulation were isolated by micromanipulation and subjected to our qPCR workflow (Fig. 2E). Based on our QC criteria, we excluded two single cells from further examination leaving twenty-eight single-cell WTA products for qPCR (S6 Table). In line with results of previous experiments, FACS and qPCR-based analyses provided concordant results. Quantitative PCR analysis enabled the detection of distinct mRNA levels of the *ERBB2* gene in MCF-10A, ZR-75-1 and MDA-MB-453 cells. In all but one comparison, i.e. MCF-10A vs ZR-75-1 (cell lines exhibiting medium and low expression levels of the *ERBB2* gene), we detected significantly different expression levels of *ERBB2* (Fig 3C). We concluded that quantification of moderate to high expression levels is feasible with our approach, while measurements are less accurate in samples exhibiting extremely low expression levels of the target gene. This may be due to noisy expression profiles (i.e. varying expression levels across different cells), suboptimal conversion rates of transcripts to WTA amplicons, technical issues related to quantification of low-abundance transcripts (24, 35, 36) or it reflects biological differences associated with low and high abundant transcripts during translation. Nonetheless, our method enabled to detect distinct levels in transcript expression in single-cell WTA products of cell lines exhibiting a 2.3-fold difference in expression of the corresponding protein. Notably, when considering all five tested breast cancer cell lines (MCF-7, MCF-10A, ZR-75-1, MDA-MB-453 and BT-474), MFI values obtained by FACS analysis highly correlated with transcript copy numbers (Pearson R = 0.925; p = 0.0243; Fig 3D).

**Fig 3.**
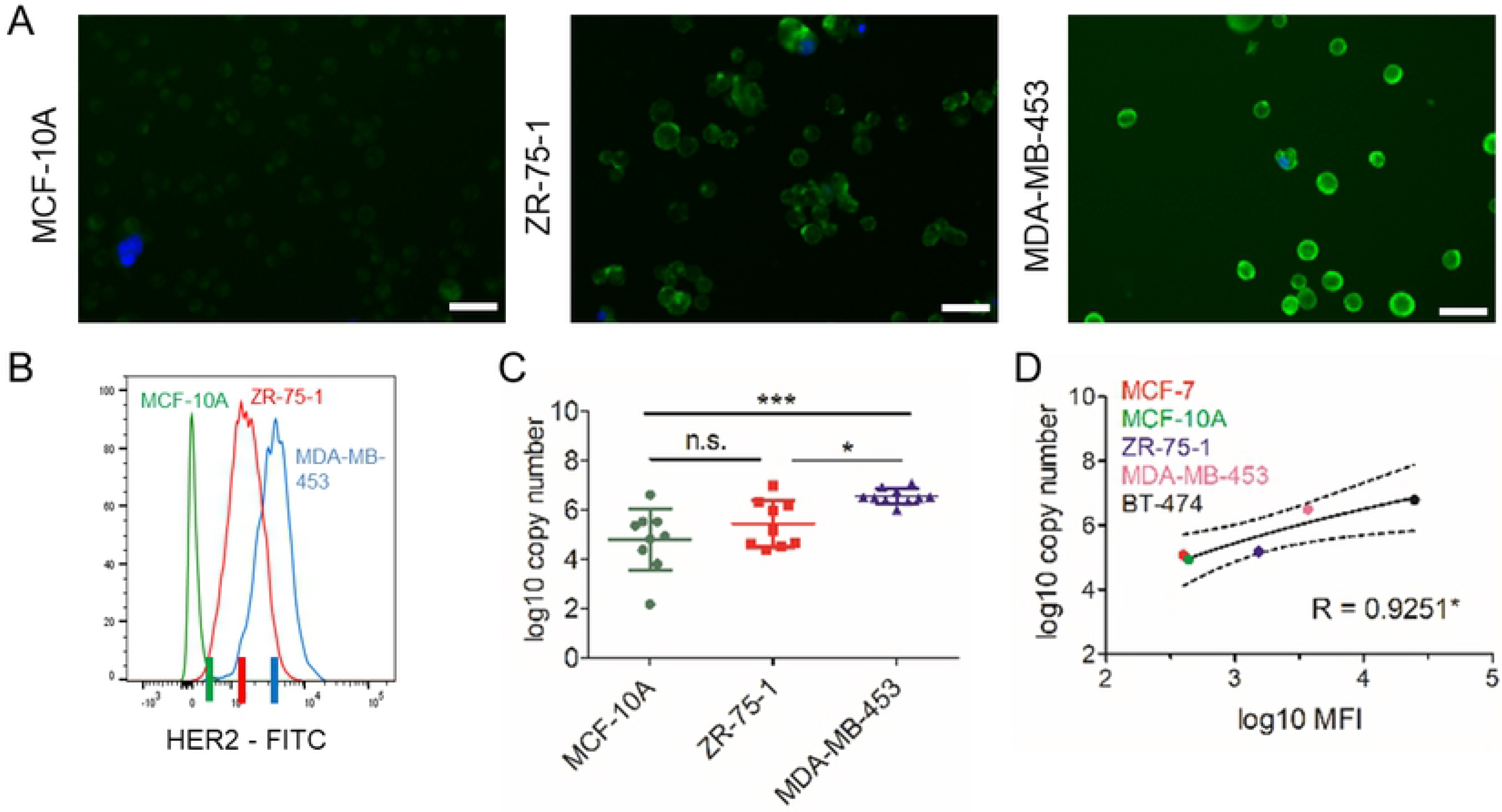
Highly accurate analysis of differential expression facilitated by the new qPCR assay. HER2 expression analysis in three breast cancer cell lines (MCF-10A, ZR-75-1 and MDA-MB-453) assessed microscopically (A) and by flow cytometry (B). (A) Brightness: +20%, contrast: −40%. Scale bar indicates 20 μm. (B) Median FITC (MCF-10A) = 436; Median FITC (ZR-75-1) = 1,529; Median FITC (MDA-MB-453) = 3,575; 3.5-fold (MCF-10A vs. ZR-75-1) and 2.3-fold (ZR-75-1 vs. MDA-MB-453) increase in HER2 expression levels. Colored lines on the *x*-axis indicate signal intensities used as thresholds for sorting of cell populations prior to single-cell isolation and analysis. (C) Quantification of *ERBB2* expression at the single-cell level was conducted following the newly established protocol (Fig. 2E). Cp values were converted to log10 copy numbers using an external standard curve. Mean ± SD; Tukey’s multiple comparisons test was applied, n.s. = not significant, * p<0.05, ** p<0.001. (D) Correlation between measured HER2 protein and *ERBB2* gene expression levels. Log10 converted median fluorescent intensity (MFI) values derived from FACS analysis and log10 converted copy numbers calculated using the absolute quantification method are plotted. Pearson’s correlation coefficient R, * p<0.05.

### Transcript quantification using re-amplified WTA products

Since the amount of cDNA contained in a single-cell WTA product is limited thus restricting the amount of downstream analyses, we sought for ways to increase the amount of the available material without introducing any considerable bias into the sample’s representation. For this reason, we developed two re-amplification protocols for WTA products differing in the sequence of used primers. Both WTA re-amplification methods employ PCR primers consisting of an universal adapter sequence identical to the one used in the primary WTA (9) at the 5’-end and a run of either fifteen or nine cytosines at the 3’-end (denoted CP2-15C and CP2-9C, respectively; S2 Table, S7-S9 Tables). Poly-C tails enable binding to oligo-G motifs present in the flanks of every WTA amplicon allowing re-amplification of primary WTA products. First, we investigated the qualitative representation of the transcriptome after re-amplification and compared fragment size distributions of PCR amplicons in the corresponding primary and re-amplified WTA products. Analysis revealed that utilization of the CP2-15C primer during WTA re-amplification resulted in an enrichment of short WTA amplicons while re-amplified WTA products generated using the CP2-9C primer showed a more similar fragment size distribution to the corresponding primary WTA products (Fig 4A and 4B). Next, we tested the impact of the annealing temperature used in the WTA re-amplification protocol (using the CP2-9C primer) on the fragment size distribution of the resulting products. Across the entire range of annealing temperatures tested (40-60°C), the fragment size distribution of re-amplified WTA products remained unchanged and therefore comparable to the primary products (Fig 4A). To examine the application of re-amplified WTA samples in qPCR, we tested both WTA re-amplification protocols (utilizing either the CP2-15C or the CP2-9C primer) using previously generated primary WTA samples from all five breast cancer cell lines. We analyzed the resulting WTA re-amplification products using the same qPCR workflow we had applied to the corresponding primary WTA samples (see above). First, we assessed the correlation between gene expression measurements obtained for primary and re-amplified WTA samples. Direct comparison of results obtained by both relative and absolute quantification methods showed high correlation between sample types (Fig 4C and 4D, S1 Fig). Quantitative PCR results obtained using both WTA re-amplification protocols (CP2-9C or CP2-15C primer) showed similar levels of correlation to the data generated with primary WTA products (Pearson R for CP2-15C: 0.98, p<0.0001 and CP2-9C: 0.96, p<0.0001; Fig 4C and 4D). Relative and absolute quantification enabled us to distinguish different expression levels of the *ERBB2* gene in re-amplified WTA products of single breast cancer cells with a comparable level of confidence as in the original WTA samples (Fig 4E, S1 Fig). Moreover, in line with the results obtained for primary WTA products, absolute qPCR quantification using re-amplified WTA samples was more accurate when conducted with purified samples as compared to specimens subjected to dscDNA synthesis (S1B Fig and S1C Fig). In summary, qPCR analysis of single-cell primary and re-amplified WTA products provided highly concordant results. Since the re-amplification protocol utilizing the CP2-9C primer was less prone to introduce a representation bias into the re-amplified WTA products, we decided to further use this variant of the protocol.

**Fig 4.**
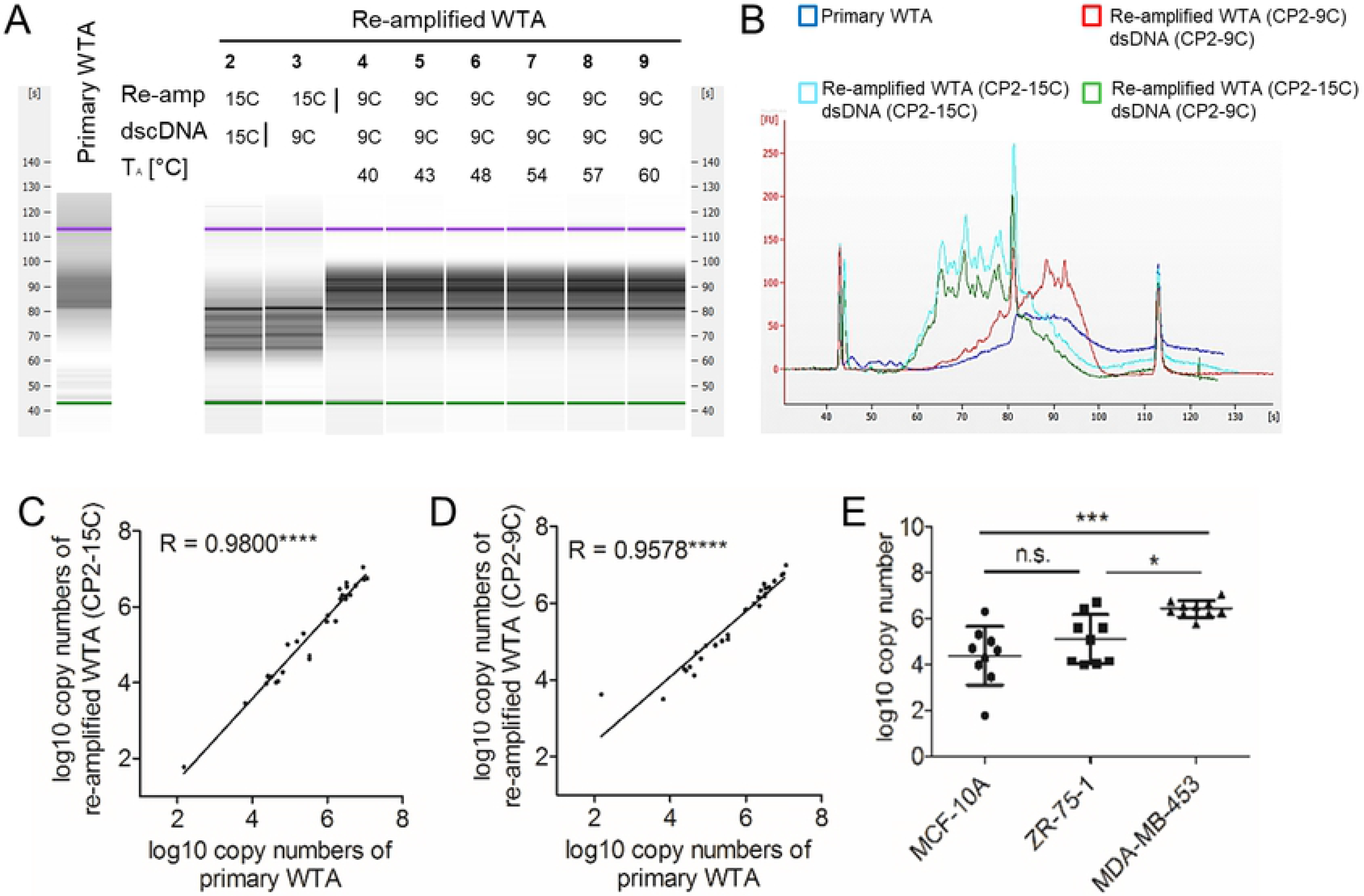
Single-cell qPCR analysis utilizing re-amplified WTA products. (A,B) Fragment size distribution of primary WTA and corresponding WTA re-amplification products (generated with either the CP2-15C or the CP2-9C primer) of one representative BT-474 single cell assessed by Bioanalyzer assay. (A) 2-9: Re-amplified WTA products generated using the indicated primer sequence for re-amplification and dscDNA-synthesis. 4-9: Samples re-amplified with different annealing temperatures (T_A_). (B) Electropherogram of selected samples. (C,D) Correlation between qPCR results conducted with primary and re-amplified WTA using CP2-15C (C) and CP2-9C (D) primers for re-amplification. (E) Absolute quantification of *ERBB2* transcript levels in MCF-10A, ZR-75-1 and MDA-MB-453 cells using re-amplified single-cell WTA products (generated using the CP2-9C primer).

### *ERBB2* expression in single disseminated breast cancer cells

Having established a workflow for qPCR-based gene expression analysis using re-amplified single-cell WTA products (Fig 2E), we tested it using patient-derived DCCs. For this, we isolated DCCs from a pleural effusion of a metastatic breast cancer patient, who had received her regular dose of the HER2-targeting antibodies trastuzumab / pertuzumab about six hours before sampling. Following a double-staining against EpCAM and HER2, single and double-positive single cells were detected and isolated by micromanipulation (Fig 5A). In total, we isolated fifteen EpCAM+/HER2− cells (i.e. EpCAM-positive DCCs lacking HER2 expression), four EpCAM^lo/−^HER2+ cells (putative DCCs low positive or negative for EpCAM and positive for HER2). The ratio of HER2^+^ / HER2^−^ cells is fully consistent with the administration of very high doses of Her-targeting antibodies few hours before ex vivo staining that successfully blocked the microscopic HER2 protein detection in the majority of cells. In addition, three double-negative cells (presumably non-epithelial, non-tumor cells) and two pools of BM cells were isolated. The isolated single cells were subjected to WTA and quality assessment as described above (Fig 2E). All but one single-cell WTA product displayed expression of at least two out of three tested housekeeping genes, meeting our QC criteria, and were included in further analyses. All re-amplified single-cell WTA products were tested positive for at least two housekeeping genes in a subsequent endpoint control PCR (S10 Table). Thereafter, all re-amplified single-cell WTA products were further processed and tested for expression of the *ERBB2* gene by endpoint PCR (Fig 5B) and qPCR (Fig 5C). Quantitative PCR analysis revealed that all HER2+ cells expressed high levels of *ERBB2* transcripts, confirming the reliability of our measurement. Interestingly, HER2− cells displayed two levels of transcript abundance: one group with a similar expression level as the HER2+ cells and another group with highly reduced or absent expression (p<0.0001, Fig 5D, S10 Table). While transcript detection in cells for which no HER2-staining could be achieved is expected after blocking the antibody-antigen binding by excessively high doses given for therapy, the detection of a population without protein and transcript detection may indicate the formation of a treatment resistant subclone.

**Fig 5.**
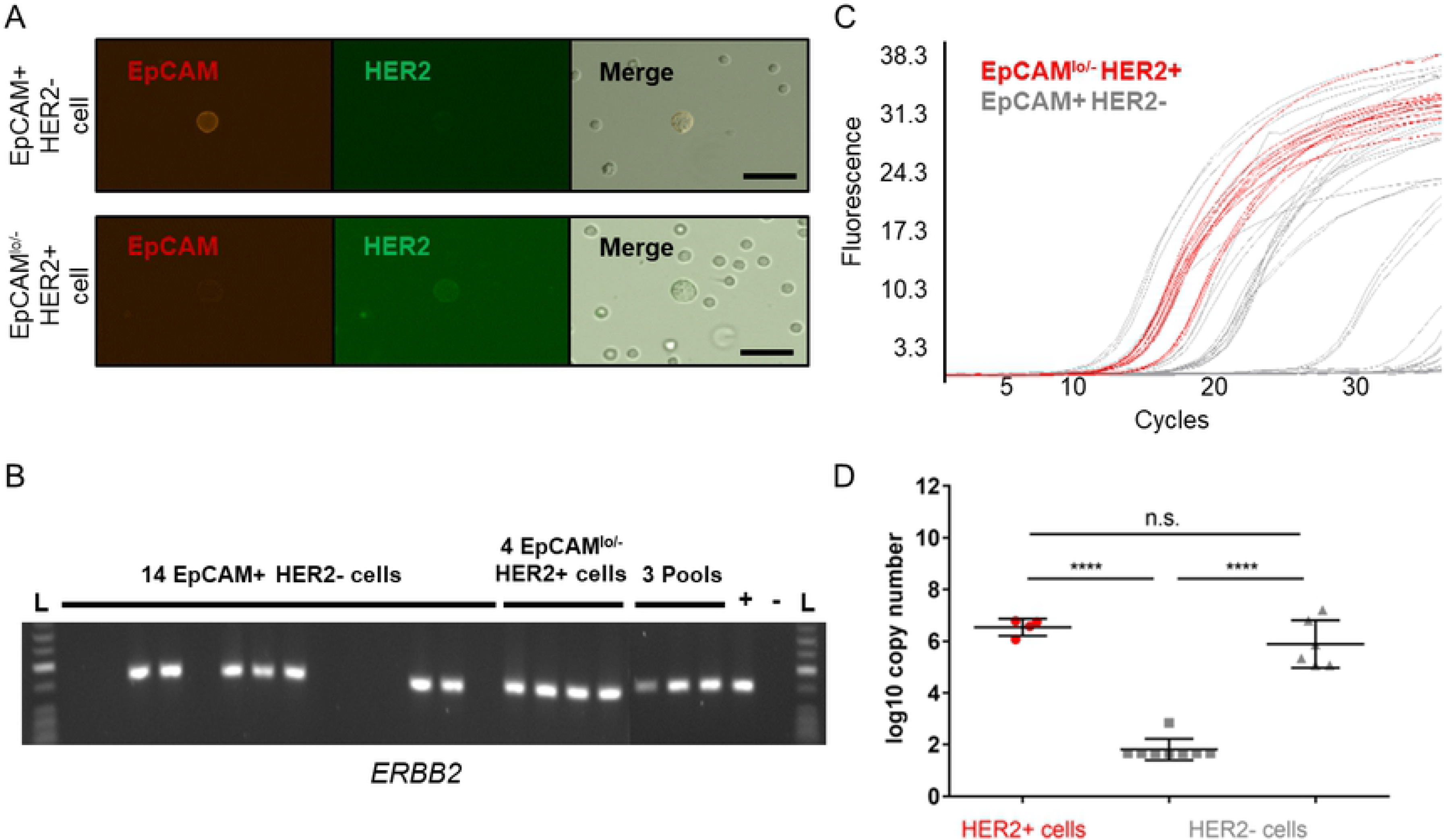
HER2/*ERBB2* expression in single cells derived from a clinical patient. (A) Cells from a pleural effusion of a metastatic breast cancer patient with a HER2-positive tumor were stained using anti-EPCAM-Cy3 and anti-HER2-FITC antibodies. Cells showing a clear membranous staining pattern were picked. Scale bar indicates 20 μm. (B) *ERBB2* expression was examined in primary WTA products by endpoint PCR. (C,D) Quantitative PCR analysis of ERBB2 expression was performed using re-amplified cDNA. (C) Amplification curves obtained for HER2+ and HER2− cells were labeled with red and gray colors, respectively. (D) Quantification of *ERBB2* expression at the single-cell level was conducted following the newly established protocol (Fig 2E). Cp values were converted to log10 copy numbers using an external standard curve. Mean ± SD; one-way ANOVA testing with Tukey’s multiple comparisons test. Color code as in C, **** p<0.0001.

## Discussion

Analyses of disseminated cancer cells (DCCs) and other rare cell populations face several unique challenges. First, DCCs are extremely rare and genetically heterogeneous making every sample precious and unique. Therefore, the analysis of such cells necessitates highly reliable single-cell technologies enabling to minimize rates of technical drop-outs and allowing highly sensitive measurements. For this reason, we established a novel workflow to reliably quantify transcript levels of selected genes using single-cell cDNA libraries generated using a previously described WTA protocol (9). The new protocol has been established to measure expression levels of the *ERBB2* gene that is of high clinical relevance. The new method enabled to measure distinct levels of *ERBB2* gene expression in cancer cell lines and patient-derived DCCs and allows addressing specific research questions with little resources.

Optimized single-cell qPCR-based protocols are very reliable, rarely suffer from technical failures (27) and exhibit very low technical noise thus allowing accurate quantification of transcripts down to ~100 copies per cell (36). For these reasons, we decided to use qPCR as read-out in our approach. To maximize the reliability of the new protocol, we carefully selected and optimized all steps of our workflow.

A key consideration in the context of qPCR-based gene expression analysis in single cells is data processing and normalization. Among many proposed strategies such as normalization to total RNA (37), cell size (38) or introduced RNA/DNA spikes (39), the most common way of normalization is to quantify the expression ratio of the target gene relative to mRNA levels of reference genes – herein referred to as relative quantification method (40). However, due to the ubiquitous cell-to-cell variability in expression of traditional reference genes, relative quantification of single cells is prone to introduce random error into single-cell analyses (41) unlike relative quantitation using bulk material (41). Various studies showed that gene expression occurs in temporal bursts also affecting housekeeping genes used as references. In relative quantification, these bursts result in false variations in transcript levels between individual cells (42–45). Indeed, our data support these reports showing considerable intra-cellular variability in gene expression of the candidate reference genes rendering most of them unsuitable for relative quantification. We noted considerable variation between the tested candidate reference genes (detection rates in single cells ranged from 11% to 93%) consistent with the reported dynamic changes in transcriptomes of single cells (46–48). To minimize the amount of missing expression data, we decided to use only the most consistently and stably expressed reference genes (*RAB7A*, *EMC7*, *REEP5*, *POLR2A*, *HPRT1*) in our analysis. Employment of these genes enabled clear discrimination between two breast cancer cell lines BT-474 and MCF-7 with high and low expression levels, respectively, of the target gene *ERBB2*. Still, despite our extensive survey for best references, exclusion of samples from data analysis due to lacking expression could not be avoided thus challenging the use of relative expression quantification for the analysis of rare cells such as DCCs.

To overcome the limitations associated with reference genes, we established a workflow utilizing absolute quantification, which requires quantification of the target gene only but necessitates careful normalization of cDNA input. For this reason, we sought for methods allowing reliable quantitation of cDNA yields in single-cell WTA products. Spectrophotometric cDNA quantification preceded by purification of WTA products proved to be the best method in our hands allowing sensitive quantification of *ERBB2* gene expression. Consequently, we conclude that the most reliable approach for normalization and quantification of single-cell qPCR data is the absolute quantification method.

In addition, to select a method for normalization and analysis of single-cell qPCR data, we sought for ways to maximize the amount of material available from each cell. For this, we developed two versions of a protocol for re-amplification of WTA products. However, we observed that WTA re-amplification is less reliable and generates high amounts of truncated amplicons when the amplification primer comprised long (15 bp long) oligo(C) motifs. Similar observations have been made when using primers containing long oligo(T) motifs during reverse transcription (49). It is likely that in both cases truncation of transcripts resulted from internal priming of cDNA amplicons. Nevertheless, a second version of the WTA re-amplification protocol, utilizing primers with shorter (9 bp long) oligo(C) priming motifs, facilitated efficient amplification of primary WTA products and reliable quantification of *ERBB2* levels by qPCR. Therefore, our new assay allows quantification of gene expression in primary and re-amplified WTA products with equal accuracy.

We tested our new workflow for its sensitivity and accuracy. The method allowed detection of both evident discrepancies in gene expression level of the *ERBB2* gene between MCF-7 and BT-474 cells as well as more subtle differences measured between MCF-10A, ZR-75-1 and MDA-MB-453 cells. In all analyses conducted in breast cancer cells, we found an excellent correlation between *ERBB2* transcript abundance and HER2 protein levels corresponding to the previously published data (33). Notably, only six cells (five in the first and one in the second comparison) had to be excluded from the analyses due to undetectable expression of *ERBB2*. This demonstrates excellent performance and accuracy of our method.

Finally, in a proof of principle experiment, we applied our new workflow on DCCs from pleural effusion fluid of a metastatic breast cancer patient with HER2-positive tumor. We categorized the collected DCCs into HER2-expressing and non-expressing groups and assessed *ERBB2* expression levels by qPCR. In agreement with the HER2 protein expression assessed by immunostaining, the expression of *ERBB2* measured by single-cell Quantitative PCR was high in the HER2+ DCCs group. Strikingly, in HER2-group (i.e. immunofluorescent negative cells for the HER2 channel), we identified single cells either expressing *ERBB2* transcripts similarly to the HER2+ cells (6/14) or highly reduced expression (8/14). Since the patient had been treated with the HER2-targeting drugs trastuzumab and pertuzumab few hours before sampling, blocking of the staining antibody was expected. In fact, the majority of detected and isolated cells were HER2− upon staining, supporting the efficient binding of the treatment antibodies. The absence of *ERBB2* transcripts may then reflect either the reported biological variation between single cells resulting from stochastic expression bursts (45) or indicate the formation of a drug-resistant clone that successfully lost addiction to the oncogene *ERBB2*. However, the faithful capture of *ERBB2* transcripts in all HER2+ cells indicate the reliability of our method in a clinical setting, whereas the detection of a transcript negative cell population may furthermore pave the way for studies addressing mechanisms of therapy resistance and clonal selection directly in patients.

## Acknowledgments

We thank Sandra Grunewald for excellent technical assistance. This work was supported by grants from the Bavarian Research Foundation (Bayerische Forschungsstiftung, DOK-165-13) and the ERC (322602).

## Supporting information

**S1 Fig.**
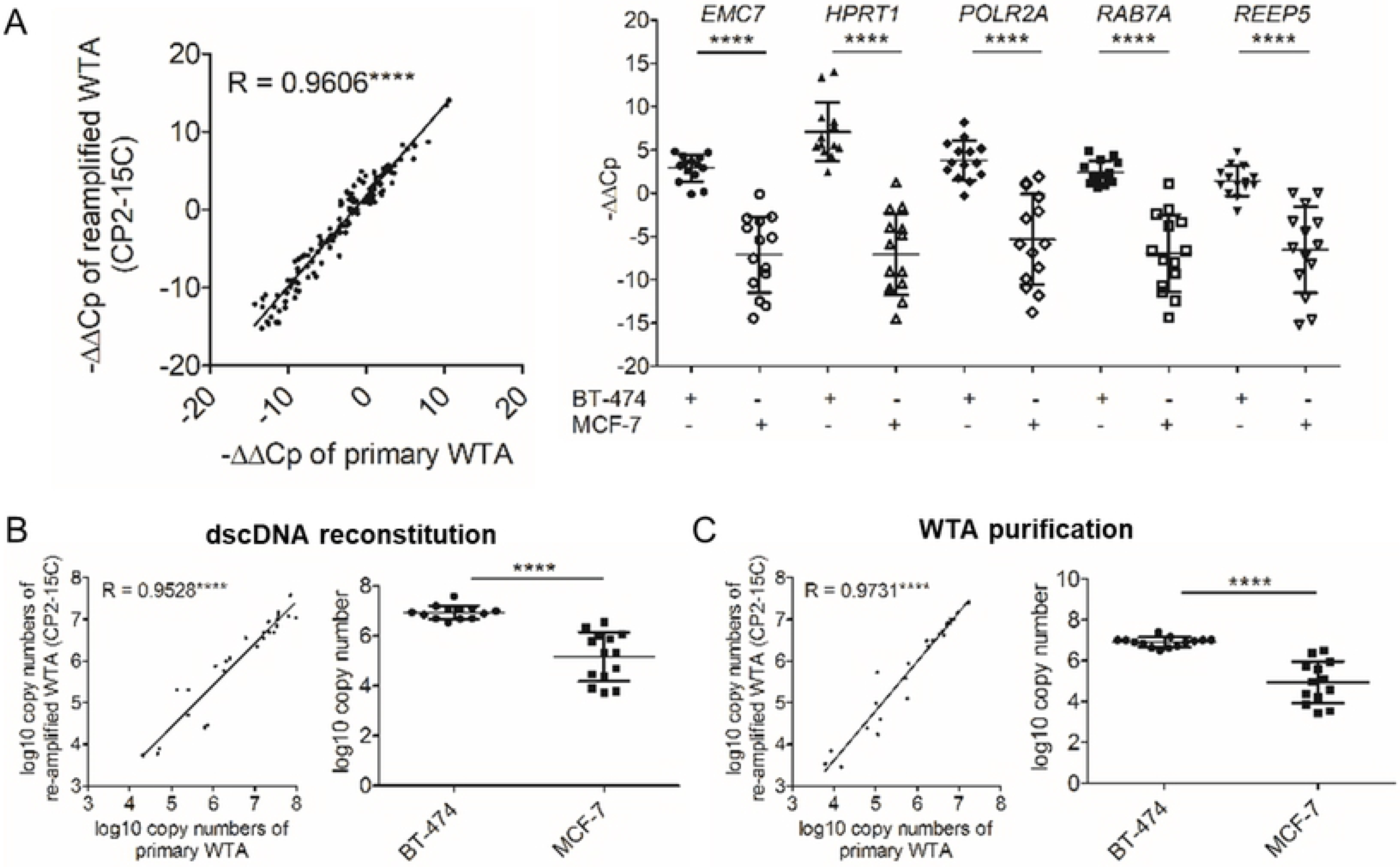
Reproducible qPCR results using re-amplified WTA products. (A) Relative quantification analysis. Correlation between log2-transformed ratios (−ΔΔCp) of re-amplified compared to primary WTA products (left panel). Spearman’s correlation coefficient R. Right panel shows relative quantification of *ERBB2* expression at the single-cell level within BT-474 and MCF-7 using single reference genes as indicated. −ΔΔCp were calculated for every single cell, mean ± SD. (B,C: left panels) Correlation of *ERBB2* qPCR results obtained by the absolute quantification strategy in re-amplified BT-474 and MCF-7 single cells between dscDNA reconstitution (B) or purified WTA (C) and diluted primary WTA products. Pearson’s correlation coefficients R. (B,C: right panels) Significant discrimination between BT-474 and MCF-7 cells regarding *ERBB2* gene expression levels. Cp values were converted to log10 copy numbers using an external standard curve. Mean ± SD; Unpaired t-test with Welch’s correction; **** p<0.0001.

**Supplementary Tables**

**S1 Table. Oligonucleotides used for amplification of target DNA sequences.**

**S2 Table. Oligonucleotides used for WTA and re-amplification**.

**S3 Table. Overall gene expression of reference genes across sample sets obtained by endpoint PCRs.**

**S4 Table. Stability of reference genes.**

**S5 Table. Gene expression analyses of primary WTA derived from BT-474 and MCF-7 single cells.**

**S6 Table. Gene expression analyses of primary WTA derived from MCF-10A, ZR-75-1, MDA-MB-453 single cells.**

**S7 Table. Gene expression analyses in re-amplified WTA (CP2-15C) of BT-474 and MCF-7 single cells.**

**S8 Table. Gene expression in re-amplified WTA (CP2-15C) of MCF-10A, ZR-75-1 and MDA-MB-453 single cells.**

**S9 Table. Gene expression in re-amplified WTA (CP2-9C) of MCF-10A, ZR-75-1 and MDA-MB-453 single cells.**

**S10 Table. Gene expression analyses of picked cells from a clinical sample.**

